# Global geographic patterns of sexual size dimorphism in birds: Support for a latitudinal trend?

**DOI:** 10.1101/012138

**Authors:** Nicholas R. Friedman, Vladimír Remeš

## Abstract

Sexual size dimorphism (SSD) is widespread among animals, and is a common indication of differential selection among males and females. Sexual selection theory predicts that SSD should increase as one sex competes more fiercely for access to mates, but it is unclear what effect spatial variation in ecology may have on this behavioral process. Here, we examine SSD across the class Aves in a spatial and phylogenetic framework, and test several a priori hypotheses regarding its relationship with climate. We mapped the global distribution of SSD from published descriptions of body size, distribution, and phylogenetic relationships across 2581 species of birds. We examined correlations between SSD and nine predictor variables representing a priori models of physical geography, climate, and climate variability. Our results show guarded support for a global latitudinal trend in SSD based on a weak prevalence of species with low or female-biased SSD in the North, but substantial spatial heterogeneity. While several stronger relationships were observed between SSD and climate predictors within zoogeographical regions, no global relationship emerged that was consistent across multiple methods of analysis. While we found support for a global relationship between climate and SSD, this support lacked consistency and explanatory power. Furthermore the strong phylogenetic signal and conspicuous lack of support from phylogenetically corrected analyses suggests that any such relationship in birds is likely due to the idiosyncratic histories of different lineages. In this manner, our results broadly agree with studies in other groups, leading us to conclude that the relationship between climate and SSD is at best complex. This suggests that SSD is linked to behavioral dynamics that may at a global scale be largely independent of environmental conditions.

## Introduction

Males and females often differ in their size, coloration, and behavior. Sexual size dimorphism (SSD) is particularly widespread (Andersson, 1994; Fairbairn *et al.*, 2007), and varies in magnitude from modest to extreme (e.g., males are up to 210% larger than females in the Great Bustard, *Otis tarda*), and from male-biased to female-biased (e.g., females are 117% larger than males in the Dapple-throat, *Arcanator orostruthus*, Székely *et al.*, 2007). Evolutionary biologists have long worked to explain this variation in terms of major selective forces and to identify its correlates. Theory predicts that in polygamous species, one sex should compete more fiercely for access to mates, and thus be selected to develop greater body size to increase its competitiveness in contests (Andersson, 1994). Mating system has emerged as the most robust and important correlate of SSD in both birds and mammals, with polygynous species exhibiting high SSD (Clutton-Brock *et al.*, 1977; Payne, 1984; Oakes, 1992; Webster, 1992; Owens & Hartley, 1998; Weckerly, 1998; Dunn *et al.*, 2001; Lindenfors *et al.*, 2003; Székely *et al.*, 2007; Lislevand *et al.*, 2009). But while many studies have examined sexual size dimorphism from a comparative or phylogenetic perspective, few have investigated geographic variation in this trait (Dunn *et al.*, 2001; Cardillo, 2002; Blanckenhorn *et al.*, 2006; Tamate & Maekawa, 2006).

Several hypotheses have been introduced to explain interspecific variation in mating systems, and could thus help predict global geographic variation in SSD. First, in terms of mating systems themselves, a classic argument suggests that the occurrence of polygyny should be related to environmental productivity, although the predicted direction differs between authors. One such argument is that productive environments might allow for aggregation of individuals (Verner & Wilson, 1966) and consequently for monopolization of female groups by males (Clutton-Brock, 1991; Lukas & Clutton-Brock, 2013). However, other researchers have argued that spatial clumping of resources might be more important (Jarman, 1974) and that homogeneous, highly productive environments should facilitate the occurrence of social monogamy (Emlen & Oring, 1977). From a temporal perspective, potential for polygamy is increased by increased temporal availability of mates, although not by extreme breeding synchrony (Emlen & Oring, 1977). As breeding seasons are shorter further from the equator, polygyny and SSD should be favored at temperate to higher latitudes. Moreover, higher variability of climate within the year leading to temporal clumping of resources might also lead to increased temporal availability of mates. Thus, polygyny and SSD might be predicted to be higher in more seasonal environments.

Second, in terms of the tenure of pair bonds, a classic argument suggests that more demanding environmental conditions require participation of both sexes in parental care (Trivers, 1972; Brown *et al.*, 2010; Royle *et al.*, 2012). Such demanding environmental conditions might include low productivity and low predictability. In line with this reasoning, it has been shown that environments with low predictability promote cooperation when raising the brood (Rubenstein & Lovette, 2007; Jetz & Rubenstein, 2011). Longer pair bonds, in turn, constrain the opportunity for polygamy and biased distribution of matings, and consequently can limit the evolution of SSD. Moreover, male-biased SSD in environments with low predictability can be also precluded by stronger sexual and social competition among females in these environments with resulting lower SSD (Clutton-Brock *et al.*, 2006; Rubenstein & Lovette, 2009).

Although sexual selection is a major influence on SSD, other hypotheses can be also useful for predicting global geographic trends in this trait. Body mass is a key adaptation of an organism to its environment, due to fundamental scaling of energetics with mass. Consequently, body mass often strongly varies with geography or environmental conditions (e.g., Bergmann’s rule; Blanckenhorn *et al.*, 2006). If larger size of males requires more energy for self-maintenance during both growth (Benito & Gonzáles-Solís, 2007; Jones *et al.*, 2009) and adulthood, males can suffer from higher mortality during times of shortage (Wikelski & Trillmich, 1997). Similarly, the evolution of particularly large males can be prevented in environments with stiff and chronic resource competition, e.g. on islands (Raia & Meiri, 2006). Chronic resource shortages (low productivity) or frequent ones (low predictability) in harsh environments would thus select against large males, opposing sexual selection and preventing the evolution of extensive SSD. Consequently, we suggest that harsh environments would select against divergence in size between sexes and thus lead to low SSD.

Although there have been many attempts to link mating system to ecology (Crook, 1964; Verner & Wilson, 1966; Jarman, 1974; Emlen & Oring, 1977; Owens & Bennett, 1997; Pérez-Barbería *et al.*, 2002), these were usually small-scale due to limitations in data availability (but see Lukas & Clutton-Brock, 2013). One previous study, conducted across 80 bird species, found no relationship between SSD and latitude after correcting for phylogeny (Cardillo, 2002). Since then, no one has yet exploited the well-established link between mating system and SSD to examine patterns of sexual selection in relation to geography, environment, and phylogeny at a global scale. Here, we make such an attempt on a large clade of animals by analyzing global variation in SSD in 2581 species of birds. We use data on SSD and environmental conditions, incorporating both geographic (species distributions) and phylogeny-based approaches, to test the following hypotheses. 1) SSD will be higher i) in temperate latitudes, where short breeding seasons should lead to intense competition for mates, and ii) in more seasonal environments, where breeding is temporarily clumped leading to stronger male-male competition. 2) SSD will be correlated with environmental productivity, either positively (clumping and consequent monopolization of females by males and/or freeing males from parental duties) or negatively (spread of biparental units across homogeneous, productive environment). 3) SSD will be lower in harsh (more variable, less predictable) environments, because these environments i) select against divergence in size between sexes, and ii) require parental cooperation in raising offspring and consequently lead to longer pair bonds, less biased distribution of matings, and lower competition for mates.

## Methods

### Species Distribution and Bioclimatic Data

We used distribution maps for bird species sampled in this study from the BirdLife International and NatureServe (2011) database, supplemented by those of Ridgely *et al.*, (2011). For use in all spatial analyses described here and below, we constructed a cylindrical equal-area projection grid with a cell area equivalent to 1° × 1° in QGIS (Brodzik & Knowles, 2002; QGIS Development Team, 2013). We overlaid polygonal range maps for each species onto this grid using Spatial Analysis in Macroecology (SAM; Rangel *et al.*, 2010) to generate gridded presence/absence and richness data. Grid cells were excluded if they did not include any landmasses, or if less than two species were present.

We chose eight bioclimatic and three geographic variables to represent our a priori hypotheses for causes of global variation in the magnitude of SSD. Of these, we derived altitude, temperature, precipitation, within-year variation in temperature, and within-year variation in precipitation from Hijmans *et al.* (2005), while deriving among-year variation in both temperature and precipitation from Harris *et al.* (2013). For each of these predictor variables, we used SAM to convert their raster data into gridded data for each cell in the global grid. Following an exploratory principle components analysis, we excluded the use of actual evapotranspiration and net primary productivity, as they were closely correlated with precipitation (Fig. S1). We combined the remaining nine predictor variables into four major models: 1) Geographic, 2) Average Climate, 3) Within-Year Climate Variation, 4) Among-Year Climate Variation (see Table 1). While these variables and models are in some cases correlated (Fig. S2), they describe different aspects of an area’s climate and biology that we are attempting to model in this study. For each chosen predictor variable, we used SAM to convert the raster data into gridded data for each cell.

**Table 1.**
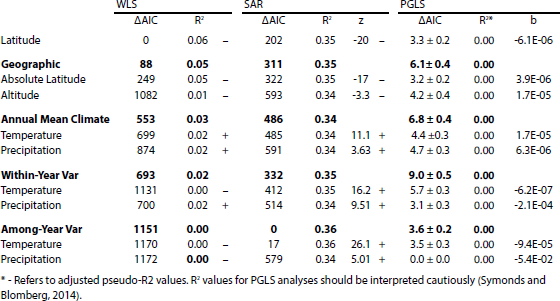
Regression of the response variable, sexual dimorphism index (SDI), as explained by environmental predictor variables. Univariate predictor regressions are shown in regular weight text, while multivariate predictor models are shown in bold with their individual component variables listed below. This table compares the goodness of fit for spatial analyses across the global grid using weighted least squares (WLS) and spatial autoregression (SAR), as well as phylogenetic comparative analyses among species using phylogenetic generalized least squares (PGLS). ΔAIC values represent the difference between each model and the best-fitting model for that analysis, with the AIC of PGLS analyses averaged across trees. The “Geographic” model includes absolute latitude but not latitude in this and all subsequent analyses.

### Sexual Size Dimorphism

We used data on body size for males and females from Lislevand *et al.* (2007). This dataset included measurements for both sexes across 2581 species that matched taxonomic descriptions in both the species distribution and phylogenetic data. We included data from each of these species in our analysis, which represents roughly a quarter of all bird species described. To describe sexual dimorphism for each species, we calculated a Sexual Dimorphism Index (SDI) following its description in Lovich & Gibbons (1992). This index is favored as a descriptor of sexual dimorphism for comparative studies (Fairbairn, 2007; Remeš & Székely, 2010). We calculated SDI as the ratio of the larger sex to the smaller sex, minus 1, making the value positive when males are larger. This differs slightly from the original SDI of Lovich & Gibbons (1992), which makes the value positive when females are larger. This was a deliberate choice on our part to make our sexual dimorphism scores intuitively interpretable as a proxy for the strength of sexual selection on males. Our reasoning is that the environmental underpinnings of polygyny are well understood, whereas those of polyandry are not (Liker *et al.*, 2013 and refs. therein). For our geographic comparisons of SDI to environmental variables, we used SAM to take the mean of the SDI values for all species present in each grid cell (Mean SDI).

### Macroecological and Phylogenetic Analyses

We used two main approaches to test for a relationship between environmental predictors and sexual size dimorphism: 1) comparison of global grid cells, each assigned the average SDI of bird species inhabiting it, and 2) comparison across species using phylogenetic comparative methods. As recommended by Blackburn & Gaston (1998), we attempted to account for variation in sampling, spatial autocorrelation, and phylogenetic non-independence. Throughout these analyses, we used an information theoretic model selection approach (Anderson, 2008), which is commonly applied to macroecological studies of this type (e.g., Jetz & Rubenstein, 2011).

For our comparison across a global grid, we used spatial autoregression (SAR) as implemented in *GeoDa* (Anselin *et al.*, 2006). Using a spatial weights grid produced with eight queen neighbors, we fit each predictor variable to Mean SDI independently, as well as a single model containing all predictor variables. We also fit weighted least squares (WLS) models to the spatial distribution of Mean SDI in *R*; these were weighted by the proportion of avian species in each grid cell for which we had data on SSD (Lislevand *et al.*, 2007; R Core Team, 2013). To assess spatial variation in the direction and strength of environmental predictors of SDI, we ran SAR and WLS analyses separately for major zoogeographical regions identified by Holt *et al.* (2013). To test for an effect of environmental conditions present only at species’ breeding sites on SSD, we repeated these spatial analyses excluding the non-breeding range of all migratory species.

To assign climatic variables to each species for our phylogenetic comparative analyses, we took the mean value of each climate variable across all grid cells occupied by a species. We also calculated the median latitude and the absolute value of the median latitude for each species. We then combined these data with a phylogeny based on the supertree of all extant birds (Jetz *et al.*, 2012), which we pruned to match our dataset. As there is considerable uncertainty relating to the topology of this supertree, we repeated the analyses described below across 100 trees sampled from the posterior distribution of the Hackett-based topology (Hackett *et al.*, 2008). To account for phylogenetic autocorrelation, we performed Phylogenetic Generalized Least Squares (PGLS) analyses on each predictor and model, as implemented under an estimated lambda model in a script accompanying Freckleton (2012), which was kindly adapted by R. Freckleton to work with the current version of the *ape* package of *R* (Version 3.0-8; Paradis *et al.*, 2004; R Core Team, 2013). We used the *geiger* package in *R* for estimations of Pagel’s lambda and disparity-through-time plots, as well as several custom scripts written in *R* available upon request (Pagel, 1999; Harmon *et al.*, 2008).

There is a widely observed relationship between body mass and SSD, which is referred to as Rensch’s rule (Rensch, 1950; see Fairbairn *et al.*, 2007). To correct for the effects of body mass on mean SDI, we repeated the analyses described above with body mass (species or grid cell average) included as a predictor variable.

## Results

### Global Patterns

Mean SDI varied considerably across the globe, with the most extreme regions ranging between −0.315 (females 31.5% larger on average) and 0.122 (males 12.2% larger). The highest degrees of SDI appear to be concentrated in areas of high breeding season productivity, such as northeastern Asia, the Neotropics, and Central Africa (Fig. 1A). Curiously however, high SDI values were also observed in regions with low biological productivity such as the Horn of Africa, the Arabian Desert, and the Sahel. Generally, SDI was negatively correlated with latitude (WLS: R^2^ = 0.06; Fig. 2A). While this linear relationship has a shallow slope, its implications are quite profound: birds in the global south are predicted to exhibit roughly twice as much male-biased SSD as birds in the global north.

**Figure. 1.**
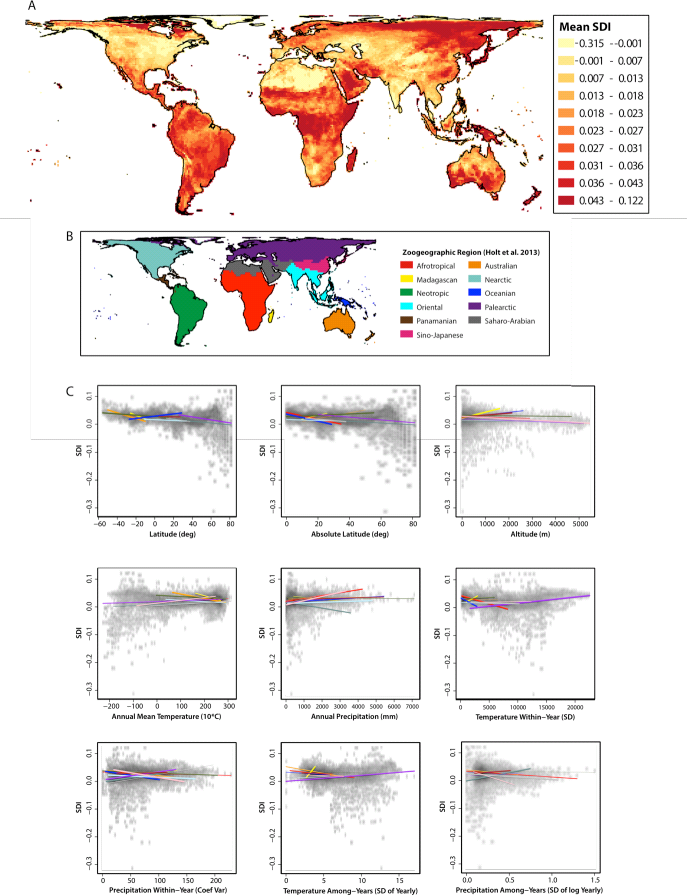
Global distribution of sexual size dimorphism and its correlates. (A) Mean Sexual Dimorphism Index (SDI) of species in each cell of a 1° equivalent equal area grid, with color classes representing 10%-ile bins. (B) Zoogeographical regions, as described by Holt *et al*. (2013). (C) Scatterplots of SDI versus the ecological predictor variables examined in this study; these have been rasterized to show point density. Trend lines represent sampling-weighted correlations for each zoogeographical region, which is labeled by color. Line width indicates model rank, with thicker lines representing more closely fitting models for that region.

**Figure. 2.**
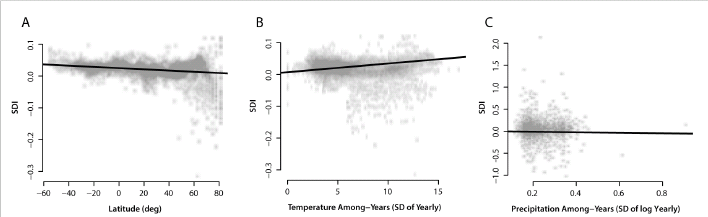
Best-fitting linear models inferred from Weighted Least Squares regression (A) across global grid cells, Spatial Auto-regression (B) across global grid cells, and Phylogenetic Generalized Least Squares (C) across species. Scatterplots have been rasterized to show point density. The Sexual Dimorphism Index (SDI) is lowest at high latitudes, and increases in areas where temperature is less predictable between years.

### Environmental Predictors

We found statistically significant correlations between all of our environmental predictors and mean SDI at the global scale; these correlations all remained significant when correcting for sampling bias and spatial autocorrelation, but not for phylogenetic non-independence (all univariate predictors were non-significant, and climate predictability was marginally non-significant; Table 1). However, there was considerable variation in the relative fit of each model depending on the type of analysis used. Among univariate models weighted for sampling bias (WLS), latitude best explained geographic variation in SDI (Table 1), with a negative relationship observed between these two variables (Fig. 1C). Among multivariate models tested, a geographic model that included absolute latitude and altitude was the strongest predictor of SDI using WLS (R^2^ = 0.05; Table 1). This suggests that birds living far from the equator or at high altitudes exhibit smaller male-biased sexual size dimorphism (Fig. 1C).

Correcting for spatial autocorrelation, among-year variation in temperature best explained geographic variation in SDI (Table 1; Figure 2B) with a positive relationship observed between these two variables. This suggests that, counter to our expectations, birds in less predictable environments may exhibit greater degrees of male-biased sexual size dimorphism. While this spatial predictor model explained a considerable amount of variation in the global distribution of SDI (with spatial lag: R^2^ = 0.35), most of this explanatory power comes from the spatial autocorrelation model (without spatial lag: R^2^ = 0.007). Correcting for average body mass did not change the rank, direction, or significance of the relationships described above (Table S1). Eliminating non-breeding portions of migratory species’ ranges resulted in apparent differences in the global distribution of SDI, particularly in the Neotropics (Fig. S3). However, correlations between breeding SDI and climate predictors behaved similarly to correlations with non-breeding ranges included, particularly in their spatial heterogeneity and strong phylogenic effect (Table S2).

Lastly, in our phylogenetic comparative analysis of variation in SDI among species, among-year variation in precipitation was the best predictor of SDI. However, this relationship was very poorly predictive (Table 1; Fig. 2C). We found that the distribution of SDI among clades strongly followed phylogenetic relationships, as indicated by our high estimate of Pagel’s lambda parameter (Pagel 1999; Figure 3). Using disparity-through-time plots (Harmon *et al.*, 2003; Figure S4), which show the accumulation of trait disparity among versus within clades, we found that variation in SDI among clades may have accumulated early in avian evolution.

**Figure. 3.**
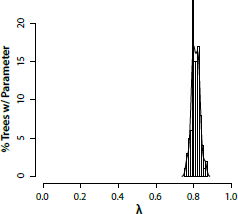
Histogram of optimum Pagel’s lambda estimates for SDI, repeated across 100 samples from the posterior tree block of Jetz *et al*. (2012). Pagel’s lambda is a tree transformation parameter whose optimum represents the amount of phylogenetic signal exhibited by a trait (Pagel 1999). A parameter value of 0 suggests weak phylogenetic signal, with the trait evolving as though on a star-tree phylogeny, while a parameter value of 1 suggests strong phylogenetic signal.

### Spatial Heterogeneity

Effects that are observed at regional scales may not be observed globally, and vice versa. To examine the relationship between climate and SSD at a finer geographic scale, we repeated our spatial analyses across the zoogeographic regions identified in Holt *et al.*, (2013). We found a considerable degree of idiosyncrasy among regions in their relationship between geography, climate, and SDI (Fig. 1C). In most regions, geographic models and within-year climate variation models best explained variation in SDI (Table S2, Table S3). Some of these correlations were quite strong, particularly those in the Afrotropic (R^2^ = 0.40), Australian (R^2^ = 0.35), and Sino-Japanese regions (R^2^ = 0.46). However, there was also considerable variation in which individual predictor variables best correlated with SDI in different regions. For example, within-year climate variation best explained variation in SDI in six out of eleven regions (Table S2), but this relationship was driven by temperature in three of these regions, and by precipitation in the other three. Similar discordance was present in the geographic models, with latitude, absolute latitude, and altitude best explaining variation in SDI within at least one region. While there was occasionally strong support for a relationship between climate and SSD, the differences in model fit among different regions and analyses demonstrate that this relationship is complex.

## Discussion

Previous studies have investigated geographical and environmental correlates of traits related to sexual selection, for example song (Botero *et al.*, 2009), plumage dichromatism (Martin *et al.*, 2010), and extra-pair paternity (Botero & Rubenstein, 2012), with only one recent study in primates focused on environmental correlates of SSD (Dunham *et al.*, 2013). Here, we conducted a global study investigating the geographic and climatic correlates of sexual size dimorphism sampling roughly one quarter of extant bird species. We identified several weak broad-scale geographical and climatic correlates of SSD in birds, but also substantial effects of geographical heterogeneity, spatial autocorrelation, and phylogeny.

Latitude was the most consistent correlate of SSD in birds both globally and within individual zoogeographic regions, with male-biased SSD increasing from the equator towards the southern pole and decreasing towards the northern pole (Fig. 1C, Tables S2, S3). This result directly contradicts our prediction that SSD should be male-biased in more seasonal northern latitudes. This result suggests a weak but measurable latitudinal gradient in SSD, but not an immediately satisfying explanation, as latitude correlates with many abiotic and biotic factors (e.g. Schemske *et al.*, 2009). Our putative explanation is that the difference between southern and northern hemisphere in SSD is the result of the unique evolutionary histories of the two avifaunas, as the effect of latitude was much weaker in phylogenetically controlled analyses (Table 1). However, this is probably not a complete explanation, because the latitudinal effect was quite consistent across zoogeographic areas (Fig. 1C), which are largely composed of evolutionarily distinct faunas (Holt *et al.*, 2013).

There remained substantial variation around global patterns of SSD in relation to climatic variables (Fig. 1C, also see Table 1), which is usually true even in smaller scale analyses (e.g., Cox *et al.*, 2003). A multitude of biotic and small-scale environmental factors were previously hypothesized or demonstrated to correlate with mating systems or SSD. They include heterogeneity in the quality of territories (Verner & Willson, 1966; Orians, 1969), population density (Lukas & Clutton-Brock, 2013), spatial and temporal clumping of resources (Emlen & Oring, 1977), prey type (Krüger, 2005; Shreeves & Field, 2008), male display behaviour (Székely *et al.*, 2000; Serrano-Meneses & Székely, 2006) or breeding in cooperative groups (Rubenstein & Lovette, 2009). It proved impossible to model effects of these factors on a global scale. However, here we were interested in global patterns and so the absence of these factors should not have biased our results.

Among zoogeographic regions, within-year variation in climate was a consistent correlate of SSD (Table S2, S3). However, while some of these local correlations were quite strong, their direction and best-fitting predictor often differed (Fig. 1C). In northern temperate regions (Nearctic and Palearctic), SDI increased with within-year climatic variability. On the contrary, in subtropical and tropical regions (Afrotropical, Oriental, and Sino-Japanese), SDI decreased with within-year climatic variability (Tables S2 and S3). One potential explanation might be that in northern temperate regions, high climatic seasonality leads to temporal clustering of available mates and thus to higher environmental potential for polygamy. On the other hand, in subtropical and tropical regions, higher environmental variability might select for cooperative breeding (Rubenstein & Lovette, 2007; Jetz & Rubenstein, 2011, but see Gonzales *et al.*, 2013), which might lead to less male-biased SSD due to high intra-sexual competition in females (Clutton-Brock *et al.*, 2006). In accordance with this hypothesis, cooperative breeding is particularly prevalent in subtropical and tropical areas (Jetz & Rubenstein 2011).

The zoogeographical regions compared in this study represent not only discrete geographical regions, but also phylogenetic clusters (Holt *et al.*, 2013). Thus, heterogeneity among these categories in their response to environmental predictors also represents heterogeneity among clades. Global correlations among climate predictors and SSD were poorly predictive after correcting for phylogeny, suggesting idiosyncratic histories for this trait in each lineage. Indeed, we found that SSD closely followed phylogeny (Figure 3), and in so doing diversified early in avian history (Figure S4). This provides some evidence that historic effects, such as biogeography and constraints on body size, may each play a major role in the evolution of SSD. Alternatively, these other relationships may be better explained by environmental filtering (Weiher & Keddy, 1999) than by correlated evolution of climatic niche and sexual size dimorphism. To disentangle the complex relationship between phylogeny and geography, ecology and evolution, studies are needed that simulate these processes to map a null distribution of species’ trait values across the globe.

In conclusion, our a priori hypotheses about global geographic and climatic correlates of SSD in birds (see Introduction) were mostly not supported. Only the hypothesis of higher male-biased SSD in regions with high climatic seasonality received partial support, and then only in northern temperate regions. Our results broadly agree with previous studies, which generally did not identify consistent climatic correlates of SSD in insects, birds, and primates (Székely *et al.*, 2004; Serrano-Meneses & Székely, 2006; Plavcan *et al.*, 2005; Dunham *et al.*, 2013; Laiolo *et al.*, 2013). There is only one study in seabirds that was able to link SSD to ocean productivity (Fairbairn & Shine, 1993), but its results remain controversial as they were contradicted by a follow-up study (Croxall, 1995). We think that all these studies together suggest that variation in SSD is likely driven by smaller-scale environmental processes, for example resource clumping on the scale relevant for avian territoriality (Verner & Willson, 1966). SSD is intimately linked to the behavioral dynamics of sexual selection, mating systems, and parental roles; we consider our results to suggest that these dynamics may be largely independent of environmental conditions (e.g. Kokko & Jennions, 2008).

## Acknowledgements

The authors would like to thank R. Freckleton for providing the PGLS script used in this study. This study was supported by the European Social Fund and the state budget of the Czech Republic (Project No. CZ.1.07/2.3.00/30.0041).

## Supporting Material

**Supplementary Figure 1:**
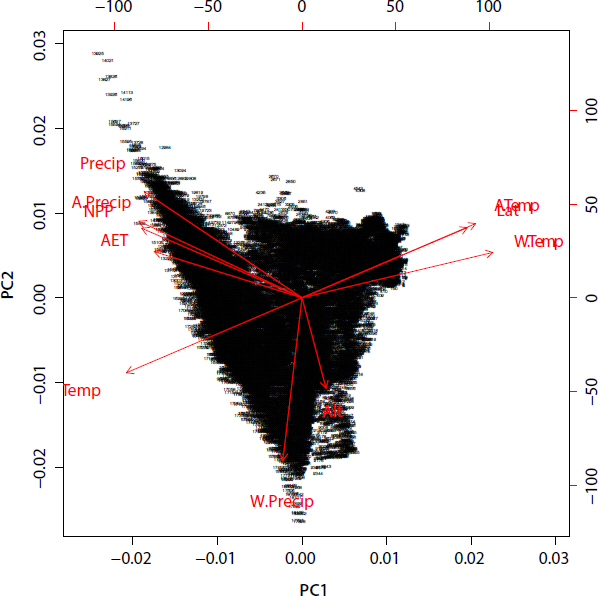
Exploratory principle components analysis of climate predictor variables. Net Primary Productivity (NPP) and Actual Evapo-Transpiration (AET) were excluded from further analyses as they closely correlate with Precipitation. The prefixes “W” and “A” refer to within-year and among-year climate variation, respectively.

**Supplementary Figure 2:**
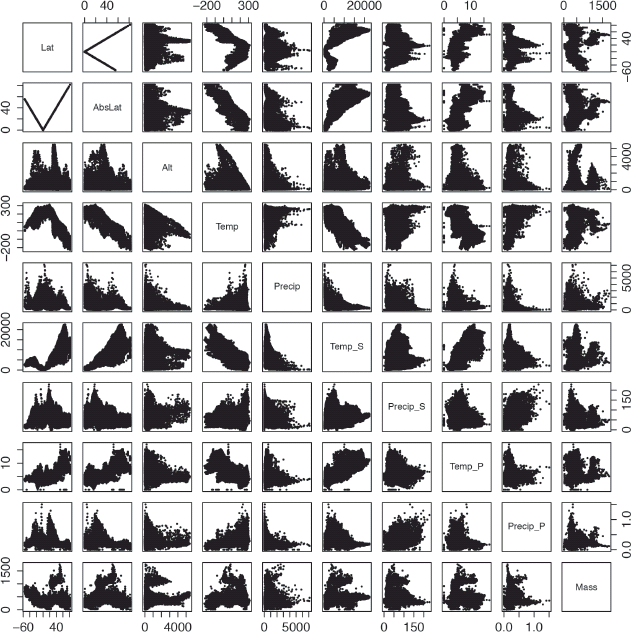
Relationships among predictor variables included in modeling global variation in SDI.

**Supplementary Figure 3:**
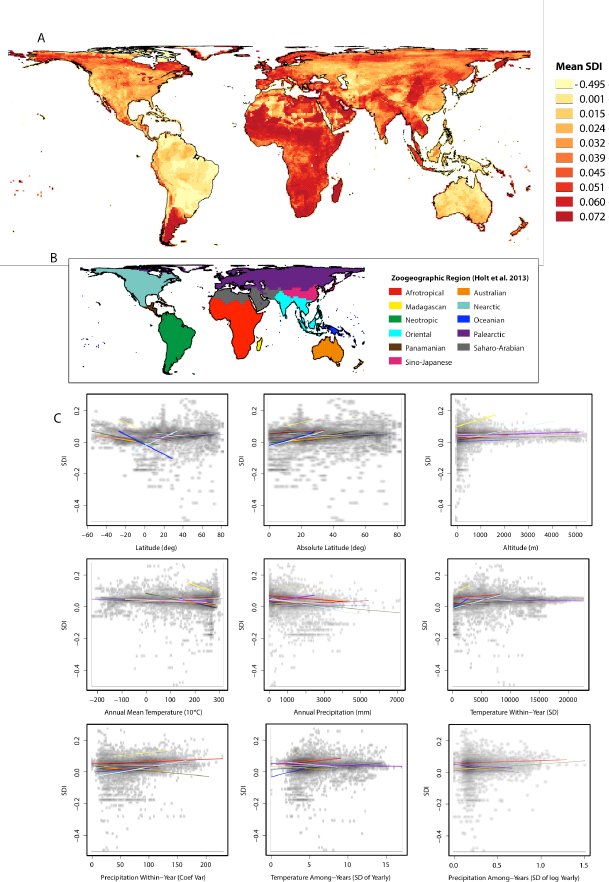
Using only breeding distributions, the global distribution of sexual size dimorphism and its climate correlates. (A) Mean Sexual Dimorphism Index (SDI) of species in each cell of a 1° equivalent equal area grid, with color classes representing 10%-ile bins. (B) Zoogeographical regions, as described by Holt et al. (2013). (C) Scatterplots of SDI versus the ecological predictor variables examined in this study; these have been rasterized to show point density. Trend lines represent sampling-weighted correlations for each zoogeographical region, which is labeled by color. Line width indicates model rank, with thicker lines representing more closely fitting models for that region.

**Supplementary Figure 4:**
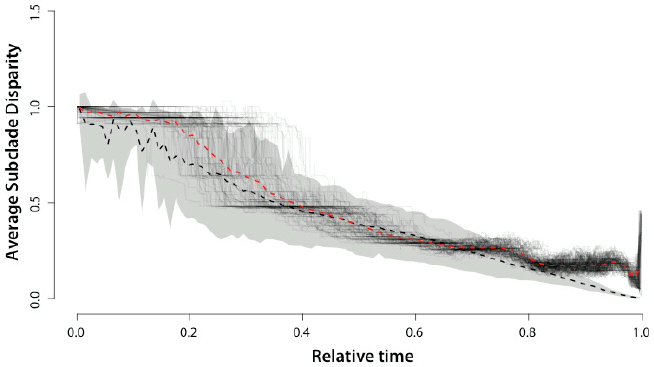
Disparity-through-time (DTT) plot of SDI for all birds. Black lines represent DTT trajectories for each of 100 trees sampled from Jetzet al. (2012), and the dotted red line is an average of these. The grey shaded area represents the 95% confidence interval of null expectation based on simulations under a Brownian motion model (see Harmon et al. 2003), and the dotted black line is an average of these. In periods with low values of subclade disparity, clades tend to occupy narrow, non-overlapping portions of character space. Thus, the early decrease in this value suggests that sexual size dimorphism diversified early among avian lineages (albeit not earlier than expected by chance).

**Table S1:** Comparison of environmental predictor models while correcting for variation in average body mass. All models are a significant fit at p < 0.001. ΔAIC values represent the difference between each model and the best-fitting model for that analysis. Multivariate models are shown in bold, with their variables listed below.

|  | WLS |  | SAR |  |  |
| --- | --- | --- | --- | --- | --- |
| | $\Delta AIC$ | | $\Delta AIC$ | z | |
| Latitude | 0 | – | 217 | -20.07 | – |
| **Geographic** | **6.6** |  | **369** |  |  |
| Absolute Latitude | 1177.93 | – | 309 | -17.48 | – |
| Altitude | 1087.64 | – | 592 | -3.63 | – |
| **Average Climate** | **523.72** |  | **483** |  |  |
| Temperature | 683.5 | + | 483 | 11.28 | + |
| Precipitation | 872.06 | + | 592 | 3.63 | + |
| **Within-Year Var** | **674.83** |  | **308** |  |  |
| Temperature | 1133.97 | – | 395 | 17.17 | + |
| Precipitation | 687.82 | + | 508 | 9.86 | + |
| **Among-Year Var** | **1157.58** |  | **0** |  |  |
| Temperature | 1177.54 | – | 19 | 26.03 | + |
| Precipitation | 1180.61 | – | 578 | 5.24 | + |

**Table S2:** Comparison of relationships between environmental predictor models and SDI calculated from species breeding ranges. Univariate predictor regressions are shown in regular weight text, while multivariate predictor models are shown in bold with their individual component variables listed below. This table compares the goodness of fit for spatial analyses across the global grid using weighted least squares (WLS) and spatial autoregression (SAR), as well as phylogenetic comparative analyses among species using phylogenetic generalized least squares (PGLS). ΔAIC values represent the difference between each model and the best-fitting model for that analysis, with the AIC of PGLS analyses averaged across trees.

|  | WLS |  | SAR |  |  | PGLS |  |  |
| --- | --- | --- | --- | --- | --- | --- | --- | --- |
| | ΔAIC | R<sup>2</sup> | ΔAIC | R<sup>2</sup> | z | ΔAIC | R<sup>2</sup>* | b |
| Latitude | 779 | 0.00 | 92 | 0.31 | 5.65 | 19.5 ± 0.8 | 0.00 | 4.20E-06 |
| **Geographic** | **606** | **0.01** |  |  |  | 13.0 ± 0.6 | **0.00** |  |
| Absolute Latitude | 847 | 0.00 | 0 | 0.31 | 11.2 | 18.1 ± 0.6 | 0.00 | 6.46E-06 |
| Altitude | 610 | 0.01 | 59 | 0.3 | 8.05 | 11.2 ± 0.7 | 0.00 | 1.81E-07 |
| **Average Climate** | **627** | **0.01** |  |  |  | **10.7 ± 0.4** | **0.00** |  |
| Temperature | 826 | 0.00 | 49 | 0.31 | -8.6 | 22.0 ± 0.6 | 0.00 | 1.27E-06 |
| Precipitation | 626 | 0.00 | 92 | 0.31 | -5.9 | 0.0 ± 0.0 | 0.00 | 2.20E-07 |
| **Within-Year Var** | **0** | **0.05** |  |  |  | **21.4 ± 0.6** | **0.00** |  |
| Temperature | 833 | 0.00 | 30 | 0.31 | 9.73 | 16.3 ± 0.5 | 0.00 | 3.10E-08 |
| Precipitation | 2 | 0.04 | 107 | 0.31 | 4.16 | 23.4 ± 0.8 | 0.00 | 3.67E-06 |
| **Among-Year Var** | **713** | **0.01** |  |  |  | **19.0 ± 0.6** | **0.00** |  |
| Temperature | 811 | 0.00 | 44 | 0.31 | 8.95 | 20.1 ± 0.6 | 0.00 | 4.92E-05 |
| Precipitation | 785 | 0.00 | 117 | 0.31 | -2.7 | 14.5 ± 0.6 | 0.00 | 1.17E-03 |

\* - Refers to adjusted pseudo-R<sup>2</sup> values. R<sup>2</sup> values for PGLS analyses should be interpreted cautiously (Symonds and Blomberg, 2014).

**Table S3:** Comparison of weighted least squared regressions across zoogeographic regions (citation). Significance level is indicated by asterices (N.S. > 0.05, * < 0.05, ** < 0.01, *** < 0.001). Direction of relationship is indicated by a + for a positive relationship, and a − for a negative relationship. Best-fit multivariate models are shaded in light grey, and best-fit predictor variables are shaded in a darker grey.

|  | Afrotropics |  |  | Australian |  |  | Madagascan |  |  | Nearctic |  |  |
| --- | --- | --- | --- | --- | --- | --- | --- | --- | --- | --- | --- | --- |
|  | ΔAIC | R<sup>2</sup> | p | ΔAIC | R<sup>2</sup> | p | ΔAIC | R<sup>2</sup> | p | ΔAIC | R<sup>2</sup> | p |
| Latitude | 1636 | 0.01 | *** - | 13 | 0.35 | *** - | 19 | 0.10 | *** - | 328 | 0.05 | *** - |
| **Geographic** | **466** | **0.31** | *** | **0** | **0.35** | *** | **0** | **0.25** | ** | **329** | **0.05** | *** |
| Absolute Latitude | 475 | 0.31 | *** - | 13 | 0.35 | *** + | 19 | 0.10 | *** + | 328 | 0.05 | *** - |
| Altitude | 1654 | 0.00 | *** - | 517 | 0.00 | N.S. - | 6 | 0.20 | *** + | 448 | 0.00 | ** + |
| **Average Climate** | **1013** | **0.18** | *** | **3** | **0.35** | *** | **9** | **0.19** | *** | **50** | **0.14** | *** |
| Temperature | 1639 | 0.01 | *** + | 10 | 0.35 | *** - | 19 | 0.10 | *** - | 376 | 0.03 | *** + |
| Precipitation | 1095 | 0.16 | *** + | 514 | 0.00 | * + | 23 | 0.06 | ** - | 238 | 0.08 | *** - |
| **Within-Year Var** | **0** | **0.40** | *** | **21** | **0.34** | *** | **5** | **0.22** | *** | **0** | **0.16** | *** |
| Temperature | 86 | 0.39 | *** - | 518 | 0.00 | N.S. + | 17 | 0.12 | *** + | 457 | 0.00 | N.S. + |
| Precipitation | 1616 | 0.02 | *** - | 82 | 0.31 | * - | 22 | 0.07 | ** + | 13 | 0.15 | *** + |
| **Among-Year Var** | **1396** | **0.08** | *** | **353** | **0.13** | ** | **1** | **0.24** | ** | **215** | **0.09** | *** |
| Temperature | 1481 | 0.06 | *** - | 359 | 0.13 | *** - | 7 | 0.19 | *** + | 449 | 0.00 | ** - |
| Precipitation | 1474 | 0.06 | *** - | 422 | 0.08 | *** - | 29 | 0.01 | N.S. + | 216 | 0.09 | *** + |
|  | Neotropic |  |  | Oceanina |  |  | Oriental |  |  | Palearctic |  |  |
|  | ΔAIC | R<sup>2</sup> | p | ΔAIC | R<sup>2</sup> | p | ΔAIC | R<sup>2</sup> | p | ΔAIC | R<sup>2</sup> | p |
| Latitude | 0 | 0.14 | *** - | 63 | 0.03 | ** + | 106 | 0.02 | *** - | 377 | 0.03 | *** - |
| **Geographic** | **33** | **0.12** | *** | **0** | **0.19** | *** | **117** | **0.01** | *** | **263** | **0.05** | *** |
| Absolute Latitude | 49 | 0.12 | *** + | 3 | 0.17 | *** - | 115 | 0.01 | *** - | 380 | 0.03 | *** - |
| Altitude | 346 | 0.01 | *** - | 64 | 0.03 | ** + | 135 | 0.00 | N.S. - | 554 | 0.00 | *** - |
| **Average Climate** | **120** | **0.09** | *** | **63** | **0.03** | N.S. | **70** | **0.05** | *** | **532** | **0.00** | *** |
| Temperature | 139 | 0.09 | *** - | 73 | 0.00 | N.S. + | 118 | 0.01 | *** - | 534 | 0.01 | *** + |
| Precipitation | 351 | 0.00 | ** - | 61 | 0.03 | *** + | 90 | 0.03 | *** + | 565 | 0.00 | *** + |
| **Within-Year Var** | **218** | **0.06** | *** | **38** | **0.10** | ** | **0** | **0.09** | *** | **0** | **0.09** | *** |
| Temperature | 297 | 0.02 | *** + | 46 | 0.07 | *** - | 120 | 0.01 | *** - | 115 | 0.08 | *** + |
| Precipitation | 254 | 0.04 | *** - | 47 | 0.07 | *** - | 16 | 0.08 | *** - | 243 | 0.06 | *** + |
| **Among-Year Var** | **357** | **0.00** | N.S. | **74** | **0.00** | N.S. | **126** | **0.01** | N.S. | **297** | **0.05** | *** |
| Temperature | 356 | 0.00 | N.S. - | 72 | 0.00 | N.S. - | 125 | 0.01 | *** + | 354 | 0.04 | *** + |
| Precipitation | 356 | 0.00 | N.S. - | 73 | 0.00 | N.S. - | 136 | 0.00 | N.S. + | 575 | 0.00 | * + |
|  | Panamanian |  |  | Saharo-Arabian |  |  | Sino-Japanese |  |  |  |  |  |
|  | ΔAIC | R<sup>2</sup> | p | ΔAIC | R<sup>2</sup> | p | ΔAIC | R<sup>2</sup> | p |  |  |  |
| Latitude | 19 | 0.00 | N.S. - | 490 | 0.02 | *** - | 454 | 0.00 | N.S. + |  |  |  |
| **Geographic** | **5** | **0.05** | N.S. | **475** | **0.03** | *** | **73** | **0.40** | N.S. |  |  |  |
| Absolute Latitude | 19 | 0.00 | N.S. - | 490 | 0.02 | *** - | 454 | 0.00 | N.S. + |  |  |  |
| Altitude | 0 | 0.04 | *** + | 508 | 0.01 | ** + | 72 | 0.40 | *** - |  |  |  |
| **Average Climate** | **0** | **0.06** | *** | **412** | **0.07** | *** | **37** | **0.43** | *** |  |  |  |
| Temperature | 19 | 0.06 | *** - | 494 | 0.01 | *** + | 152 | 0.33 | *** + |  |  |  |
| Precipitation | 16 | 0.01 | N.S. + | 497 | 0.01 | *** + | 54 | 0.41 | *** + |  |  |  |
| **Within-Year Var** | **19** | **0.00** | N.S. | **2** | **0.28** | N.S. | **0** | **0.46** | ** |  |  |  |
| Temperature | 19 | 0.00 | N.S. + | 0 | 0.00 | * - | 453 | 0.00 | N.S. + |  |  |  |
| Precipitation | 17 | 0.00 | N.S. + | 497 | 0.28 | *** + | 9 | 0.45 | *** - |  |  |  |
| **Among-Year Var** | **17** | **0.01** | N.S. | **459** | **0.04** | *** | **284** | **0.21** | *** |  |  |  |
| Temperature | 18 | 0.00 | N.S. - | 497 | 0.01 | *** + | 285 | 0.00 | N.S. - |  |  |  |
| Precipitation | 16 | 0.00 | N.S. + | 463 | 0.03 | *** - | 451 | 0.20 | *** - |  |  |  |

**Table S4:** Comparison of spatial autocorrelation regressions across zoogeographic regions (citation). Significance level is indicated by asterices (N.S. > 0.05, * < 0.05, ** < 0.01, *** < 0.001). Direction of relationship is indicated by a + for a positive relationship, and a − for a negative relationship. Best-fit multivariate models are shaded in light grey, and best-fit predictor variables are shaded in a darker grey.

|  | Afrotropics |  |  | Australian |  |  | Madagascan |  |  | Nearctic |  |  |
| --- | --- | --- | --- | --- | --- | --- | --- | --- | --- | --- | --- | --- |
|  | ΔAIC | z | p | ΔAIC | z | p | ΔAIC | z | p | ΔAIC | z | p |
| Latitude | 1548 | -22.5 | ** | 16.35 | -17 | *** | 21.32 | -2.5 | * | 401.7 | 9.7 | *** |
| **Geographic** | **377** |  |  | **1.25** |  |  | **0** |  |  | **385** |  |  |
| Absolute Latitude | 391 | -35.2 | *** | 16.35 | 17 | *** | 21.32 | 2.5 | * | 401.7 | 9.7 | *** |
| Altitude | 1540 | -4.25 | *** | 126.9 | -2.4 | * | 2.197 | 5.3 | *** | 476.1 | -4.2 | *** |
| **Average Climate** | **853** |  |  | **0** |  |  | **10.4** |  |  | **255** |  |  |
| Temperature | 1522 | 6 | *** | 1.46 | -16.7 | *** | 15.74 | -3.6 | *** | 343.2 | -12.5 | *** |
| Precipitation | 957 | 12.9 | *** | 131.8 | 0.8 | N.S. | 23.27 | -2 | * | 296.1 | -14.3 | *** |
| **Within-Year Var** | **0** |  |  | **0.8** |  |  | **7.07** |  |  | **0** |  |  |
| Temperature | 71 | -46.1 | *** | 131.1 | -1.2 | N.S. | 18.05 | 3.1 | ** | 173 | 18.5 | *** |
| Precipitation | 560 | 23.8 | *** | 29.84 | -12.2 | *** | 19.1 | 3 | ** | 263.5 | 17.8 | *** |
| **Among-Year Var** | **991** |  |  | **102** |  |  | **7.27** |  |  | **311** |  |  |
| Temperature | 1011 | 0.63 | N.S. | 101.6 | -13.2 | *** | 12.56 | 4.1 | *** | 367.3 | 11.4 | *** |
| Precipitation | 992 | -14.7 | *** | 117.2 | -4 | *** | 27.02 | 0.7 | N.S. | 440.5 | 10.8 | *** |
|  | Neotropic |  |  | Oceanian |  |  | Oriental |  |  | Palearctic |  |  |
|  | ΔAIC | z | p | ΔAIC | z | p | ΔAIC | z | p | ΔAIC | z | p |
| Latitude | 0 | -14.2 | *** | 42.67 | 0.8 | N.S. | 53 | -1.3 | N.S. | 237.4 | -16.5 | *** |
| **Geographic** | **24.2** |  |  | **14** |  |  | **52** |  |  | **206.5** |  |  |
| Absolute Latitude | 49.7 | 12.9 | *** | 14.68 | -7.3 | *** | 55 | 0 | N.S. | 239.4 | -16.5 | *** |
| Altitude | 155 | -3.6 | *** | 41.47 | 1.4 | N.S. | 51 | -2.1 | * | 486.4 | -3 | ** |
| **Average Climate** | **129.1** |  |  | **16.7** |  |  | **53** |  |  | **296.9** |  |  |
| Temperature | 128 | -7.8 | *** | 43.26 | 0.3 | N.S. | 55 | -0.3 | N.S. | 335 | 12.9 | *** |
| Precipitation | 166.4 | -1.2 | N.S. | 14.79 | 5.5 | *** | 51 | 2.1 | * | 428.8 | 8.2 | *** |
| **Within-Year Var** | **109** |  |  | **0.75** |  |  | **34** |  |  | **347.8** |  |  |
| Temperature | 147.8 | 5.1 | *** | 31.21 | -3.6 | *** | 54 | 0.7 | N.S. | 353.9 | 12 | *** |
| Precipitation | 129.6 | -6.3 | *** | 0 | -6.8 | *** | 35 | -6 | *** | 482.3 | 3.7 | *** |
| **Among-Year Var** | **168.4** |  |  | **30.1** |  |  | **0** |  |  | **0** |  |  |
| Temperature | 166.7 | -1 | N.S. | 29.11 | -3.9 | *** | 10 | 6.8 | *** | 37.1 | 22.4 | *** |
| Precipitation | 167.7 | 0.2 | N.S. | 40.74 | -1.6 | N.S. | 40 | 3.9 | *** | 481 | 3.8 | *** |
|  | Panamanian |  |  | Saharo-Arabian |  |  | Sino-Japanese |  |  |  |  |  |
|  | ΔAIC | z | p | ΔAIC | z | p | ΔAIC | z | p |  |  |  |
| Latitude | 19.18 | -1.2 | N.S. | 156.3 | 2 | * | 24.27 | -8.5 | *** |  |  |  |
| **Geographic** | **1.99** |  |  | **148** |  |  | **0** |  |  |  |  |  |
| Absolute Latitude | 19.18 | -1.2 | N.S. | 156.3 | 2 | * | 24.27 | -8.5 | *** |  |  |  |
| Altitude | 0.63 | 4.5 | *** | 150 | -3.2 | ** | 84.95 | 3.6 | *** |  |  |  |
| **Average Climate** | **0** |  |  | **148** |  |  | **63.1** |  |  |  |  |  |
| Temperature | 1.04 | -4.5 | *** | 146.3 | 3.7 | *** | 87.69 | 2.5 | * |  |  |  |
| Precipitation | 15.54 | 2.2 | * | 156.3 | -1.9 | N.S. | 65.85 | 5.6 | *** |  |  |  |
| **Within-Year Var** | **14.83** |  |  | **0** |  |  | **13.2** |  |  |  |  |  |
| Temperature | 20.42 | -0.3 | N.S. | 159.8 | -0.6 | N.S. | 87.58 | -2.2 | * |  |  |  |
| Precipitation | 12.86 | 2.8 | ** | 2.33 | 12.9 | *** | 11.63 | -10 | *** |  |  |  |
| **Among-Year Var** | **16.96** |  |  | **162** |  |  | **30.6** |  |  |  |  |  |
| Temperature | 18.69 | 1.4 | N.S. | 159.9 | -0.4 | N.S. | 92.19 | 0.1 | N.S. |  |  |  |
| Precipitation | 18.36 | 1.5 | N.S. | 160 | 0.3 | N.S. | 35.5 | -7.7 | *** |  |  |  |

